# Built to change: dominance strategy changes with life stage in a primitively eusocial bee

**DOI:** 10.1101/2020.07.30.229625

**Authors:** Margarita Orlova, Erin Treanore, Etya Amsalem

## Abstract

Access to reproduction is determined by an individual dominance’s rank in many species and is achieved through aggression and/or dominance signalling. In eusocial insects one or several dominant females (queens) monopolize reproduction but to what extent queens rely on aggression and signalling remains obscure. Aggression is costly and its efficiency depends on the group size, whereas signalling may reduce the risks and costs of aggression. Both strategies are used to regulate reproduction in social taxa, with aggression being more common in small social groups, compared to signalling in larger societies.

Here, we examine the use of aggression and chemical signalling in a social species (*Bombus impatiens*) where the dominant queen interacts with increasing numbers of workers as she ages.

We found that the queens’ strategy to monopolize reproduction changes with life stage, shifting from overt aggression to chemical signalling as the queen gets older. Particularly, old queens exhibited a higher ratio of short to long cuticular hydrocarbons compared to young queens, an endogenous shift that was attributed to age, as all egg-laying queens were fecund and kept with the same number of workers. Our findings contribute to the understanding of reproductive dominance in the context of an individual’s life history.

## Introduction

An individual’s access to reproduction is often linked to their place in the dominance hierarchy (Moxon, 2009), and dominant individuals frequently enjoy higher reproductive success than the subordinates in many taxa (Ang and Manica, 2010a; Bulger, 1993; Frank, 1986; Monnin and Peeters, 1999). In eusocial animals, where reproduction is monopolized by the dominant individual/s and the subordinate workers function as helpers (Hamilton, 1964; Reeve and Keller, 2001), the ability to achieve dominance is critical for individual fitness.

The most common strategy to attain a high rank in the dominance hierarchy is via aggressive behaviour (Clarke and Faulkes, 2001; Monnin and Peeters, 1999). While effective for maintaining reproductive dominance, aggression is costly and may have adverse effects on the fitness of the aggressor (De Luca and Ginsberg, 2001; Gobin et al., 2003). In some social animals, the group is so large that fighting with every subordinate would be impractical. Aggression in such societies is sometimes exhibited by hopeful reproductives (future queens), vying for higher status among themselves (Gilley, 2001; Monnin et al., 2003), but not against non-reproductive subordinates (workers) who heavily outnumber the dominant (the queen). In these species, the dominant surpasses the subordinates in its reproductive capacity rather than in fighting ability, and aggressive interactions between the queen and the workers occur very rarely and are frequently lethal for the queen (DeGrandi-Hoffman et al., 2007; Orlova et al., 2019). Maintenance of reproductive monopoly of the dominant under such conditions often depends on signalling.

Dominance signalling is an alternative strategy used to maintain reproductive dominance while avoiding the costs of fighting (le Roux and Bergman, 2012; Rohwer, 1975). Signalling can be costly as well, even though the costs are much harder to estimate given our limited knowledge of the physiological mechanisms of signal production. Signals may be limited by the physical properties of the medium in which they operate, e.g. lighting, temperature (Evans and Norris, 1996), by its own physicality (e.g., chemistry, volatility) and by the evolution of communication strategies in the species of interest (Endler et al., 1993). However, the main limitation of signalling as a dominance strategy lies in the fact that its efficiency depends on both the receiver and the signaller. Dominance signalling can only be an evolutionarily stable strategy if it is honest and advantageous for both the signaller and the receiver (Smith and Harper, 1995). The honesty of dominance signalling is ensured either by its prohibitively high cost or by its inextricable link to specific physiological traits (Smith, 1994; Zahavi, 1975). In social animals, the honesty of signals is especially important since in case of dishonest dominance signals produced by the queen, the sterile workers would suffer significant fitness losses (Keller and Nonacs, 1993).

The efficiency of aggression and signalling as means to achieve high dominance status is heavily dependent on the social environment. For example, group size and aggressiveness of group members limit the efficiency of aggression as a dominance strategy, so that in very large or very aggressive groups the dominant has less opportunities to monopolize resources including reproduction (Amsalem and Hefetz, 2011; Ang and Manica, 2010b; Balasubramaniam et al., 2012). On the other hand, response to dominance signalling can be influenced by the responder’s own qualities and situational context (Martín et al., 2007; Tibbetts, 2008). Thus, for an individual living in a changing social environment, flexible use of each of these strategies would be optimal. We hypothesize that in a species where the change in one of the parameters of social environment, e.g. group size, is an integral part of the species biology, the dominant individual would exhibit plasticity in using both strategies.

To test this, we focused on a species where the social environment changes significantly in the course of the dominant individual’s lifetime, the bumble bee *Bombus impatiens*, and examined the use of aggression and chemical signalling by the dominant individual to regulate subordinate’s reproduction at different life stages, representing a change in the size of the social group.

In bumble bees the queen initiates a colony as a solitary individual, and during her life cycle, the colony size gradually increases reaching up to several hundreds of individuals. Thus, the young founding queen interacts with just a few workers but as the colony develops, she is confronted with larger numbers of individuals that can potentially challenge her reproductive dominance. Aggression plays an important role in the determination of reproductive dominance both among workers and between the queen and the workers (Amsalem and Hefetz, 2010; Padilla et al., 2016), and its efficacy depends on group size (Bloch and Hefetz, 1999b; Van Doorn, 1989). Other studies suggested a significant role for chemical signalling (Roseler et al., 1981; Van Oystaeyen et al., 2014), but these claims have been questioned (Amsalem et al., 2015; Bloch and Hefetz, 1999a). We decided to tackle both the behaviour and the chemical signalling simultaneously to better understand by what means *Bombus impatiens* queens maintain their reproductive dominance at different stages in their lives. To do so, we grouped queens of different life stages (unmated, newly mated, young and old egg-laying queens) with two workers and examined the aggressive behaviour exhibited by and towards the queens and among the workers. Additionally, we non-lethally sampled the cuticular hydrocarbons profile of these queens repeatedly and further tested the inhibitory effect of these queens on worker reproduction. Finally, we constructed a model to examine whether aggressive behaviour and chemical signalling at each life stage are predictors of workers reproduction.

## Methods

### Bees

Colonies of *B. impatiens* (n=8) were obtained from Koppert Biological Systems (Howell Michigan, USA), maintained in the laboratory under constant darkness, a temperature of 28–30°C, 60% relative humidity, and supplied *ad libitum* with a 60% sugar solution and fresh pollen (Light spring bee pollen, 911Honey). These colonies were used as a source of newly emerged workers (younger than 24 h) and queens.

### Experimental design

Newly-emerged workers were collected upon emergence, paint-marked and housed with an unrelated queen in a plastic cage (11 cm x 7 cm) for 7 days. Four types of queens were used in the experiment: (1) unmated, (2) newly mated, (3) young egg-laying and (4) old egg-laying queens (see below). The cages were observed for 10 minutes daily for 7 days. Aggressive behaviours performed by all individuals were recorded. All queens underwent a non-lethal sampling of cuticular hydrocarbons (hence CHCs) as described below on days 0 (before being housed with workers), 1, 3, 5 and 7 of the experiment. On day 7, all workers, unmated and newly mated queens were frozen. Young and old egg-laying queens were housed with a new pair of newly-emerged workers and observed for another 7 days. Thus, CHCs of young and old egg-laying queens were sampled every other day for 14 days.

### Queen types

Four types of queens were generated as follows. Newly-emerged, unmated queens were collected from Koppert colonies and grouped with workers upon emergence. A portion of the newly-emerged unmated queens were aged for 6 days, mated in the laboratory with unrelated males and underwent CO_2_ treatment to mimic the diapause required for colony foundation according to the protocol developed by (Amsalem and Grozinger, 2017) and described below. Newly-mated queens were sampled for CHCs 24 hours following the CO_2_ treatment and then housed with workers. A portion of the mated queens were housed in individual cages until they laid eggs and produced workers. These young egg-laying queens were housed with newly-emerged workers within 7 days from the emergence of their first daughter. Old egg-laying queens were obtained directly from Koppert colonies several months following the emergence of the first worker. These colonies contained >100 workers and were producing gynes and males. Overall, we used 35 queens and 94 workers in this study.

### CO_2_ treatment

Individual queens were placed in a closed (but not entirely sealed) cage. A tube connected to a pure CO_2_ tank was inserted into the cage and a stream of CO_2_ was blown into the cage for 60 seconds. It was visually ascertained that the queens were completely anaesthetized within approximately 20 seconds. Following this treatment, each queen was kept in the same cage for 30 minutes in the rearing room until the anaesthesia wore off, which typically happened within 20 minutes. The cages were then ventilated and queens were transferred to new cages.

### Behavioural observations

Observations were carried out daily under regular light. Cages of an unknown treatment to the observer were observed in a random order. Aggressive behaviours of the queen and workers (which were individually marked with a color tag) were recorded daily for 10 minutes per cage. We recorded the identity of the individual at which a particular behaviour was directed and constructed indices of behaviours performed by the queen toward workers (hence “aggression performed by the queen”), performed by workers towards the queen (hence “aggression received by the queen”) and performed among workers (hence “aggression among workers”). Aggressive interactions included climbing (one bee mounting another bee), humming (rapid wing movements directed at another bee without a physical contact), darting (rapid movement towards another bee without a physical contact), pushing (physical contact from which the other bee retreats) and attack (overt fight with biting and stinging attempts), as described in (Amsalem and Grozinger, 2017; Amsalem and Hefetz, 2010). All these behaviours are performed in a higher rate by dominant bumble bee females, both workers and queens (Amsalem and Grozinger, 2017; Amsalem and Hefetz, 2010; Amsalem and Hefetz, 2011; Amsalem et al., 2014a; Amsalem et al., 2014b; Duchateau, 1989; Padilla et al., 2016). The sum of all aggressive behaviours performed by the queen, received by the queen or among workers that occurred during the observation period per cage was used in further analyses.

### Non-lethal sampling of CHCs

In order to continuously sample the same queen every other day, we designed a protocol for non-lethal sampling of the queen cuticular profile. Each queen was placed in an individual clean 20 ml scintillation glass vial for 10 minutes, and then placed back to her cage. The vial was then washed with 1 ml hexane and the solution was transferred into a clean 2 ml glass vial. The solution was then evaporated to 200 µl and transferred to a clean insert placed in the same vial. The solution was then nearly fully evaporated and was washed with 50 µl hexane containing 2 µg pentadecane (Sigma) as an internal standard. The resulting extract was evaporated to 20 µl, of which 1 µl containing 100 ng of internal standard was further analysed using GC/MS.

### Measurement of ovarian activation

All workers were 7 days old upon freezing. Workers were dissected under a stereomicroscope. Ovaries were obtained and placed into drops of distilled water. The length of the terminal oocyte in the three largest ovarioles was measured with a micrometer eyepiece embedded into the lens. Workers possess four ovarioles per ovary and at least one oocyte per ovary was measured. Mean terminal oocyte length for each bee was used as an index of ovarian activation (Amsalem et al., 2009).

### Chemical analysis

To identify the compounds in the queen cuticular profile we pooled individual samples from queens of each type and analysed them using an Agilent 7890A GC equipped with a HP-5ms column (0.25id x 30m x 0.25µm film thickness, Agilent Technologies, Santa Clara, CA) connected to an Agilent 5975C mass spectrometer. The run was performed in splitless mode with temperature program from 60 °C at 15 °C/min for 4 min to 120 °C, then at 4 °C/min for 54 min to 300 °C with a final hold of 5 min. The resulting chromatograms and spectra were analysed using MSD ChemStation software (Agilent) and all peaks were identified using the NIST database and by comparing retention times and mass signatures with synthetic compound standards. All individual samples were then quantified with gas chromatograph Trace 1310 (ThermoScientific) equipped with a TG-5ms column. The run was performed in splitless mode with a temperature program as above. All chromatograms were integrated using Chromeleon 7.0 software (ThermoScientific). Compounds were identified based on diagnostic ions in the resulting spectra, and by matching retention times and spectra with authentic standards. Peak areas were normalized to the internal standards and are presented as percentage of the total secretion to account for differences between queens.

### Statistical analysis

Statistical analyses were performed using SPSS v.21. Generalized Estimating Equations analysis (hence GEE) was employed for comparisons of worker ovary activation, aggression levels and percentage of compounds between queens. The models were built to control for interdependencies within data using queen identity as a subject variable. Time at sampling was used as a within-subject variable. Unstructured correlation matrix was used in models for oocyte size and aggressive behaviour. Robust estimation was used to handle violations of model assumptions. In all analyses we used queen type as the main effect followed by post-hoc contrast estimation using Least Significant Difference (LSD) method. Discriminant analysis was used to compare chemical profiles in their entirety between queen types. Generalized Linear Mixed Model analysis was performed to assess the contribution of different factors to workers reproductive suppression. Since young and old egg-laying queens were observed for two consecutive weeks, the models were built to control for interdependencies within data using ‘queen identity’ as a subject variable and ‘week’ as a within subject variable. Unstructured correlation matrix was used in all models. Generalized Linear Mixed Model analysis was performed on standardized values (Z-scores) to obtain standardized beta coefficients. Data of oocyte size, chemical parameters and aggressive behaviour are presented as boxplots featuring the minimum and maximum values, outliers and medians. Statistical significance was accepted at α=0.05.

## Results

Comparison of the average terminal oocyte of workers that were kept with different queen types revealed that workers housed with old egg-laying queens had the least activated ovaries, while those housed with unmated queens had the most activated ovaries. Ovary activation of workers housed with mated and young egg-laying queens were intermediate (GEE, Wald χ^2^_3_ = 10.34, p=0.016; Figure 1).

**Fig. 1.**
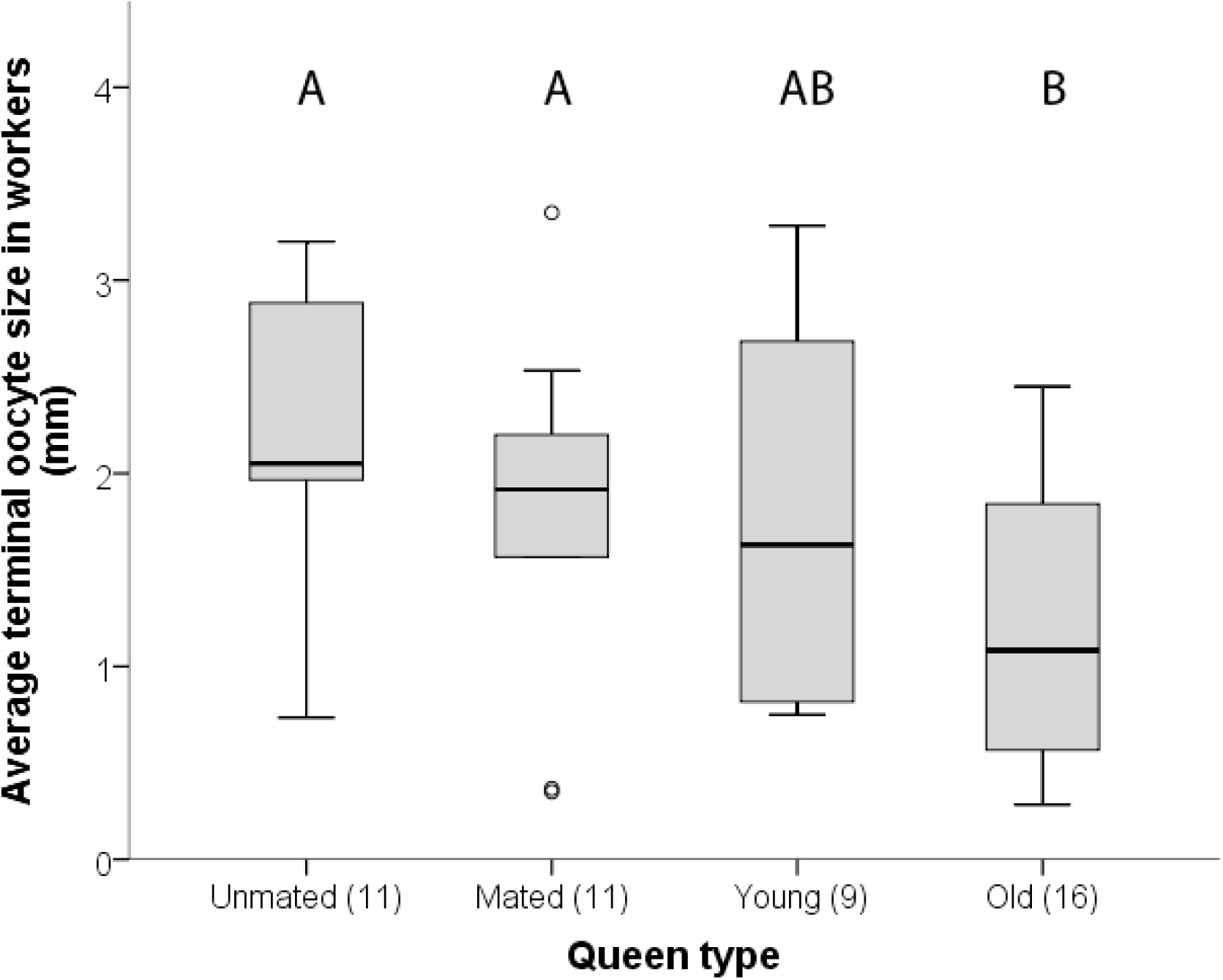
Average terminal oocyte size in worker pairs headed by queens of different type. Box plots display medians, quartiles and minimum and maximum values. Dots above/below each box indicate outliers. Sample sizes are indicated in parentheses, significant differences are indicated by different letters above columns

Aggressive behaviour differed significantly between worker pairs grouped with different queens. Both aggression performed by queens towards workers and by workers towards queens were lower in pairs grouped with old egg-laying queens compared to all other queen types (GEE, Wald χ^2^_3_ = 18.91, p<0.001 and GEE, Wald χ^2^_3_ = 38.79, p<0.001 for aggression performed and received by the queen, respectively). Aggressive behaviours performed or received by the queen or among workers did not differ between unmated, mated, and young egg-laying queens (Fig. 2). Aggressive behaviours among workers were lower in pairs grouped with old egg-laying queens compared to unmated and young egg-laying queens. In pairs grouped with mated queens, aggression levels among workers were intermediate (GEE, Wald χ^2^_3_ = 12.28, p=0.006; Fig 2).

**Fig. 2.**
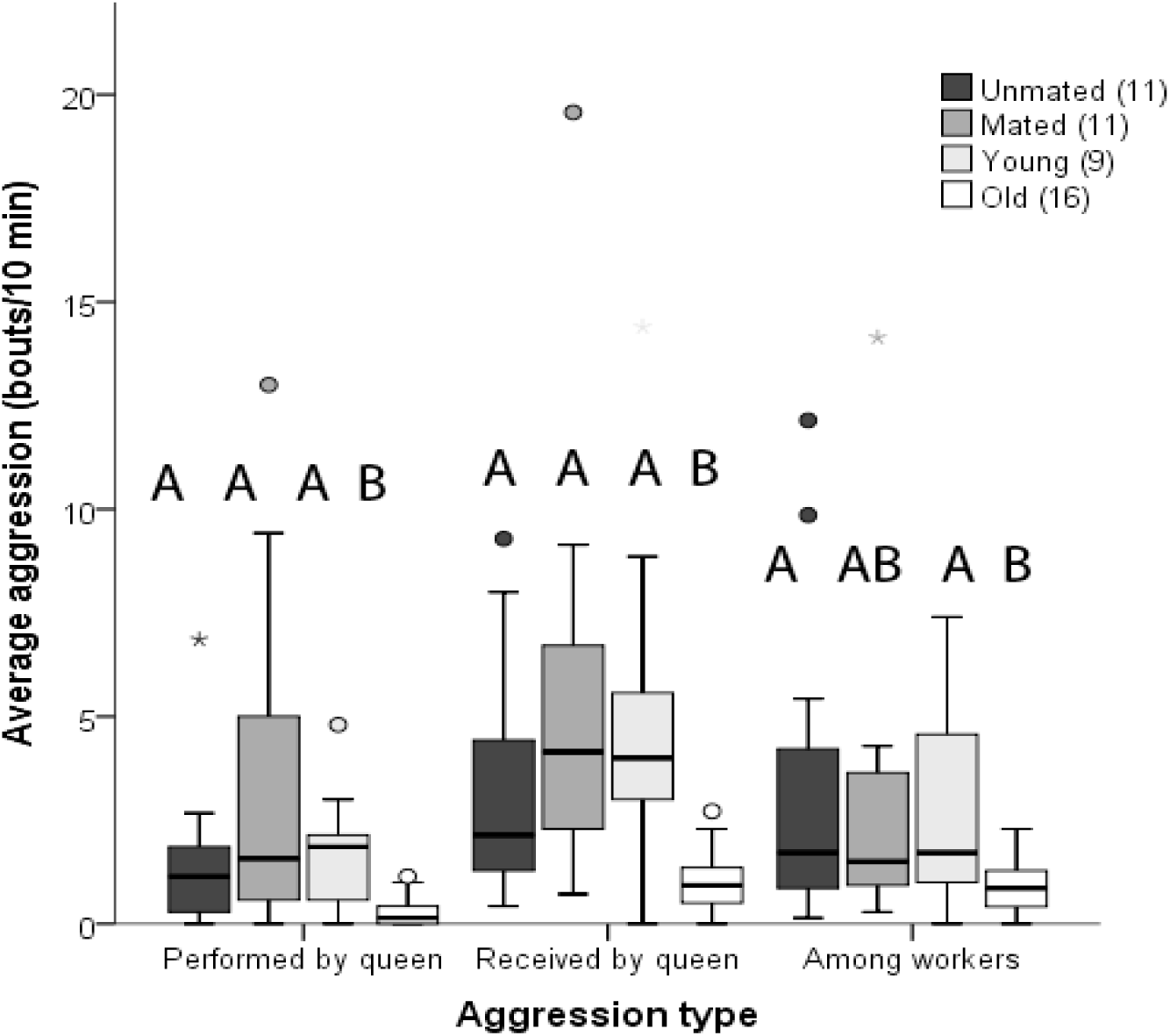
Aggressive behaviour in groups headed by queens of different type. Box plots display medians, quartiles and minimum and maximum values. Dots above/below each box indicate outliers. Different queen types are represented by different colours. Sample sizes are indicated in parentheses, significant differences are indicated by different letters above columns.

A total of 23 components were identified in the cuticular profile of all queen types and were produced by all queens. These included alkanes, alkenes, long-chain acetate esters, fatty alcohols and aldehydes (Table 1). Cuticular profiles of queens were analysed using discriminant analysis using relative quantities of cuticular substances. Three discriminant functions significantly discriminated between queen types using this analysis, with the first two functions explaining 91.6% of the variance. Function 1 (eigenvalue = 4.59, canonical correlation = 0.91, percent of explained variance = 73.6%; Wilk’s λ_66_ = 0.06, χ^2^ = 56.89, p<0.001) discriminated between old egg-laying queens and all other queens and had highest correlation values with major hydrocarbon components, while function 2 (eigenvalue= 1.32, canonical correlation = 0.73, percent of explained variance = 18%; Wilk’s λ_42_ = 0.31, χ^2^ = 23.64, p<0.001), discriminated between unmated, mated and young egg-laying queens and had highest correlation values with acetate and aldehyde components (Fig. 3).

**Table 1.** Percentages of cuticular compounds (from the total secretion) in different queen types (presented as mean ± s.e.m).

|  |  | **Mated** | **Old** | **Unmated** | **Young** |
| --- | --- | --- | --- | --- | --- |
| **Peak #** | **Peak Name** | **Mean (%)**<br>**±SEM** | **Mean (%)**<br>**±SEM** | **Mean (%)**<br>**±SEM** | **Mean (%)**<br>**±SEM** |
| **1** | Hexadecanal | 0.34±0.06 | 0.26±0.1 | 0.76±0.2 | 0.1±0.02 |
| **2** | Octadecanal | 0.25±0.05 | 0.08±0.02 | 0.57±0.15 | 0.06±0.02 |
| **3** | Octadecanol | 0.35±0.1 | 0.12±0.03 | 0.3±0.1 | 0.23±0.03 |
| **4** | heneicosane | 1.72±0.23 | 0.8±0.04 | 1.53±0.21 | 0.75±0.09 |
| **5** | docosane | 0.27±0.02 | 0.37±0.02 | 0.23±0.01 | 0.27±0.02 |
| **6** | Eicosenol | 0.55±0.28 | 0.3±0.23 | 0.62±0.21 | 0.08±0.02 |
| **7** | Tricosene | 7.26±0.63 | 10.39±1.15 | 6.19±0.78 | 6.69±1.05 |
| **8** | Tricosane | 16.07±0.52 | 27.11±1.12 | 16.47±0.94 | 17.44±1.2 |
| **9** | Tetracosene | 0.31±0.03 | 0.63±0.06 | 0.34±0.04 | 0.41±0.07 |
| **10** | Tetracosane | 0.91±0.33 | 1.47±0.34 | 0.71±0.05 | 0.77±0.05 |
| **11** | Pentacosene | 14.9±0.89 | 16.2±1.23 | 15.69±1.13 | 13.79±1.12 |
| **12** | Pentacosane | 16.2±0.76 | 21.7±0.94 | 19.89±1.24 | 17.67±0.71 |
| **13** | Hexacosene | 0.24±0.01 | 0.11±0.01 | 0.28±0.02 | 0.2±0.02 |
| **14** | Hexacosane | 0.49±0.02 | 0.27±0.03 | 0.53±0.03 | 0.47±0.05 |
| **15** | Heptacosene | 8.34±0.53 | 1.93±0.15 | 6.66±0.34 | 5.9±0.88 |
| **16** | Heptacosane | 10.02±0.53 | 4.32±0.2 | 11.85±0.84 | 8.62±0.92 |
| **17** | Tetracosyl acetate | 4.23±0.4 | 6.25±0.72 | 2.88±0.51 | 9.54±1.59 |
| **18** | Nonacosene | 5.57±0.42 | 1.11±0.08 | 3.98±0.26 | 4.11±0.59 |
| **19** | Nonacosane | 4.21±0.32 | 2.05±0.16 | 4.74±0.35 | 3.16±0.38 |
| **20** | Hexacosyl acetate | 3.95±0.28 | 2.87±0.3 | 2.98±0.69 | 6.26±0.75 |
| **21** | Hentriacontene | 2.56±0.25 | 0.98±0.09 | 1.32±0.16 | 2.38±0.34 |
| **22** | Hentriacontane | 0.94±0.14 | 0.42±0.04 | 1.11±0.18 | 0.48±0.14 |
| **23** | Octacosyl acetate | 0.31±0.03 | 0.26±0.02 | 0.37±0.04 | 0.59±0.06 |

**Fig. 3.**
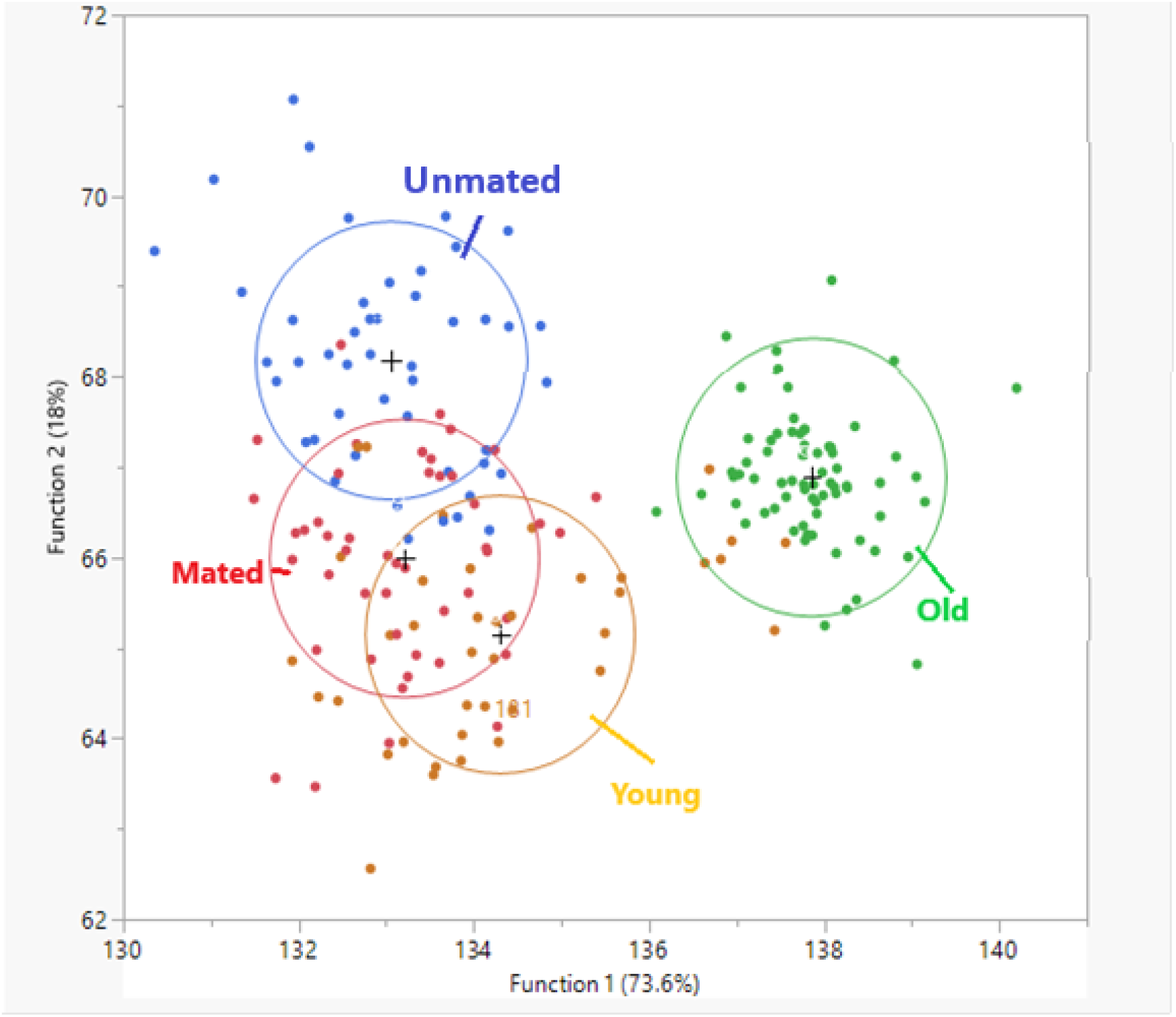
Difference in cuticular profile between different queen types. Different queen types are indicated by different colours, ellipses contain 50% of samples, variance explained by each function is indicated in parentheses.

Since most of the variance between queen types (73.6%) was explained by the hydrocarbon profile (function 1), ten compounds with the absolute largest correlation coefficients with function 1 were used for further analysis. These compounds included docosane (C22), tricosene (C23:1), tricosane (C23), tetracosene (C24:1), hexacosene (C26:1), hexacosane (C26), heptacosene (C27:1), heptacosane (C27), nonacosene (C29:1) and nonacosane (C29). Examination of correlation coefficients revealed that the longer-chained hydrocarbons (equal or longer than 26 carbons), were negatively correlated with function 1, while the shorter-chained ones (equal or shorter to 24 carbons) were positively correlated with it. We termed the former “long CHCs” and the latter “short CHCs” and calculated the ratio between the short and the long (S:L) CHC ratio to represent the chemical composition of the cuticle in further analysis. Pentacosane (C25), previously identified as a queen pheromone in *B. terrestris* (Van Oystaeyen et al., 2014) was not included in the 10 most influencing hydrocarbons in the queen cuticle throughout her life cycle. Its levels were highest in unmated and old egg-laying queens, lowest in newly-mated queens and intermediate in young egg-laying queens, with no significant effect of time since the start of the experiment (GEE, Wald χ^2^_3_ = 20.51, p<0.001 for queen type, Wald χ^2^_8_ = 10.35, p=0.237 for day of sampling).

S:L CHC ratio differed significantly across queen types and days from the onset of the experiment, and there was significant interaction between queen type and time from the start of the experiment (GEE, Wald χ^2^_3_ = 53.32, p<0.001 for queen type, Wald χ^2^_8_ = 32.77, p<0.001 for day of sampling, Wald χ^2^_16_ = 145.02, p<0.001 for interaction). Post-hoc analysis revealed that old egg-laying queens had higher ratio of S:L CHCs compared to other queen types, but no difference was found between mated, unmated and young egg-laying queens (post-hoc LSD, p<0.001 for old egg-laying queens vs. other queen types, p>0.1 for all other comparisons; Fig. 4). S:L CHC ratio changed significantly with time in mated and young egg-laying, but not in unmated and old egg-laying queens. In unmated queens, the ratio of S:L CHCs dropped on day 1 and then increased gradually until reaching the original values on day 7 (post-hoc LSD, p<0.05 for day 1 vs. day 7, with intermediated values on days 3 and 5, data not shown). In mated queens, the S:L CHC ratio decreased with time and was much lower on day 7 than on day 0 (post-hoc LSD, p<0.005, with intermediate values on days 1, 3 and 5; data not shown). In young egg-laying queens, the ratio increased with time and was significantly higher on days 8, 10, 12 and 14 than on days 0, 1 and 3 (p<0.03 for all comparisons; Fig. 5). In old egg-laying queens, no significant differences between different time points were found (p>0.05 for all comparisons, data not shown).

**Fig. 4.**
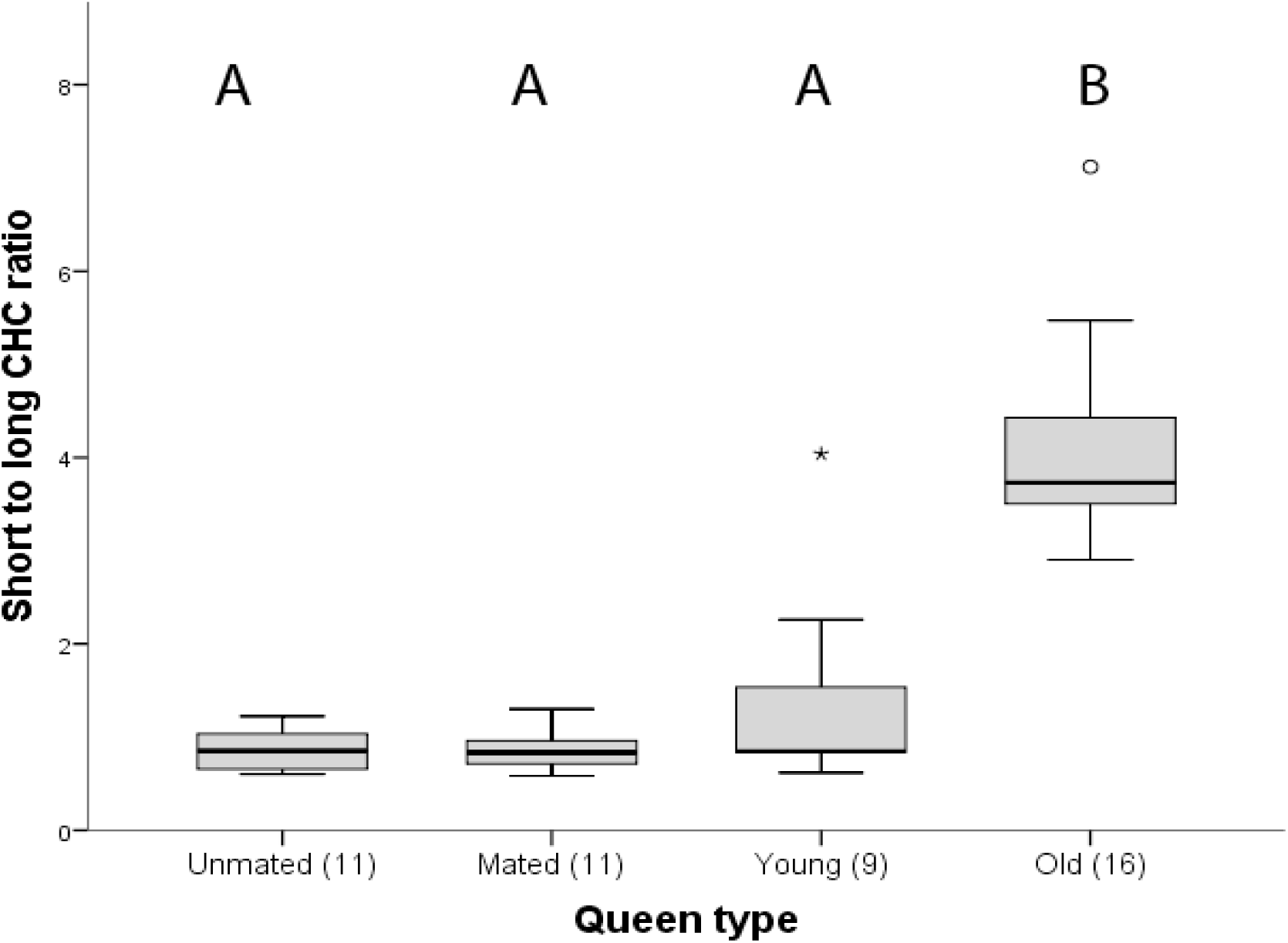
Short to long CHC ratio in queens of different type. Box plots display medians, quartiles and minimum and maximum values. Dots above/below each box indicate outliers. Sample sizes are indicated in parentheses, significant differences are indicated by different letters above columns.

**Fig. 5.**
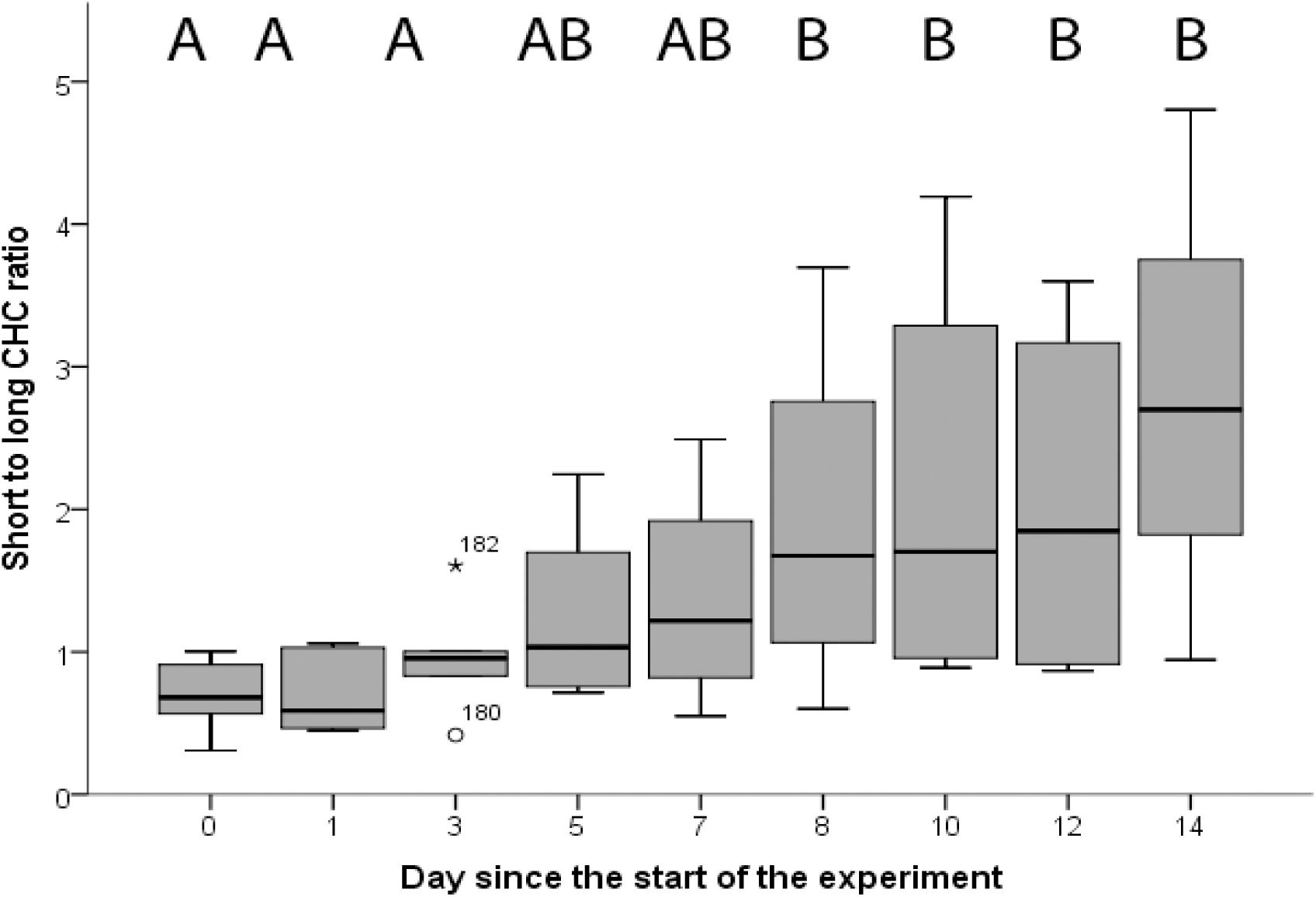
Short to long CHC ratio in young egg-laying queens (n=5) during 14 days since the onset of the experiment. Box plots display medians, quartiles and minimum and maximum values. Dots above/below each box indicate outliers. Significant differences are indicated by different letters above columns.

Aside from hydrocarbons, other compounds present on the cuticle also differed significantly between the queen types. These compounds included tetracosyl, hexacosyl and octacosyl acetates, hexadecanal and octadecanal. The three acetates were significantly positively correlated with function 2 of the discriminant analysis that explained 18% of total variance between queen types, while the two aldehydes, though present in very small amounts (see Table 1), were negatively correlated with it. Percentage of acetates was significantly higher in young egg-laying queens compared to all other queen types (GEE, Wald χ^2^_3_ = 11.626, p=0.009), while the percentage of aldehydes was highest in unmated queens and lowest in young egg-laying queens (GEE, Wald χ^2^_3_ = 13.969, p=0.003).

To examine the predictors of oocyte size in workers, we constructed several generalized linear models to assess the influence of queen aggressive behaviour and chemical signalling on worker ovarian activation. In these models, we used worker oocyte length as a dependent variable and queen type, aggression performed and received by the queen, aggression among workers and S:L CHC ratio in queens as predictor variables, as well as interaction between queen type and each continuous predictor variable. The full model had a better fit compared to any of the reduced models (ΔCAIC=10 or higher). In the full model, the effects of queen type, aggression performed and received by the queen, S:L CHC ratio and interactions of queen type with all types of aggression were significant (Table 2). We also built a model with the same structure as described above substituting short to long CHC ratio with pentacosane proportion. This model had a poorer fit (ΔCAIC=10) and pentacosane did not significantly predict worker oocyte length by itself or significantly interact with other factors.

**Table 2.** Parameters of the generalized linear mixed model of effects of behavioural and chemical factors on worker oocyte size.

| **Factor** | **F** | **df1,2** | **p-value** |
| --- | --- | --- | --- |
| **Queen type** | 38.198 | 3,10 | 0.0001 |
| **Aggression performed by the queen** | 28.896 | 1,3 | 0.019 |
| **Aggression received by the queen** | 11.12 | 1,7 | 0.012 |
| **Aggression among workers** | 0.06 | 1,3 | 0.943 |
| **CHC ratio** | 5.154 | 1,25 | 0.032 |
| **Queen type\*Aggression performed** | 12.267 | 3,25 | 0.0001 |
| **Queen type\*Aggression received** | 6.704 | 3,25 | 0.002 |
| **Queen type\*Aggression among workers** | 12.573 | 3,25 | 0.0001 |
| **Queen type\*CHC ratio** | 11.331 | 3,2 | 0.068 |

Since significant interaction was found between the queen types and all continuous predictors, we decided to build a separate model for testing the effects of predictors for young and old egg-laying queens, showing that different parameters were significant predictors of worker oocyte size in different queen types. The parameters of each model are displayed in Table 3. In young egg-laying queens, aggression performed by the queen was the only significant negative predictor for worker oocyte size. In old egg-laying queens, aggression among workers was a significant positive predictor, and S:L CHC ratio was a significant negative predictor of worker oocyte size, while the effect of aggression performed by the queen was not significant. Overall, the strongest negative predictors of worker oocyte size in queens, that were able to reduce worker reproduction, were aggressive behaviour performed by young egg-laying queens and CHC profile in old egg-laying queens (Fig. 6).

**Table 3.** Parameters of the generalized linear mixed models for effects of young and old queens respectively on worker oocyte size

|  | Young egg-laying queen |  |  |  | Old egg-laying queen |  |  |  |
| --- | --- | --- | --- | --- | --- | --- | --- | --- |
| **Factor** | **F** | **df1,2** | **p-value** | **Coefficient** | **F** | **df1,2** | **p-value** | **Coefficient** |
| **Aggression performed by the queen** | 35.86 | 1,4 | 0.005 | -2.66 | 0.537 | 1,11 | 0.47 | 0.17 |
| **Aggression received by the queen** | 3.3 | 1,4 | 0.143 | 0.96 | 0.674 | 1,11 | 0.43 | 0.39 |
| **Aggression among workers** | 0.55 | 1,4 | 0.5 | -0.4 | 26.205 | 1,11 | 0.001 | 1.78 |
| **CHC ratio** | 8.07 | 1,3 | 0.055 | -1.05 | 11.711 | 1,11 | 0.006 | -0.56 |

**Fig. 6.**
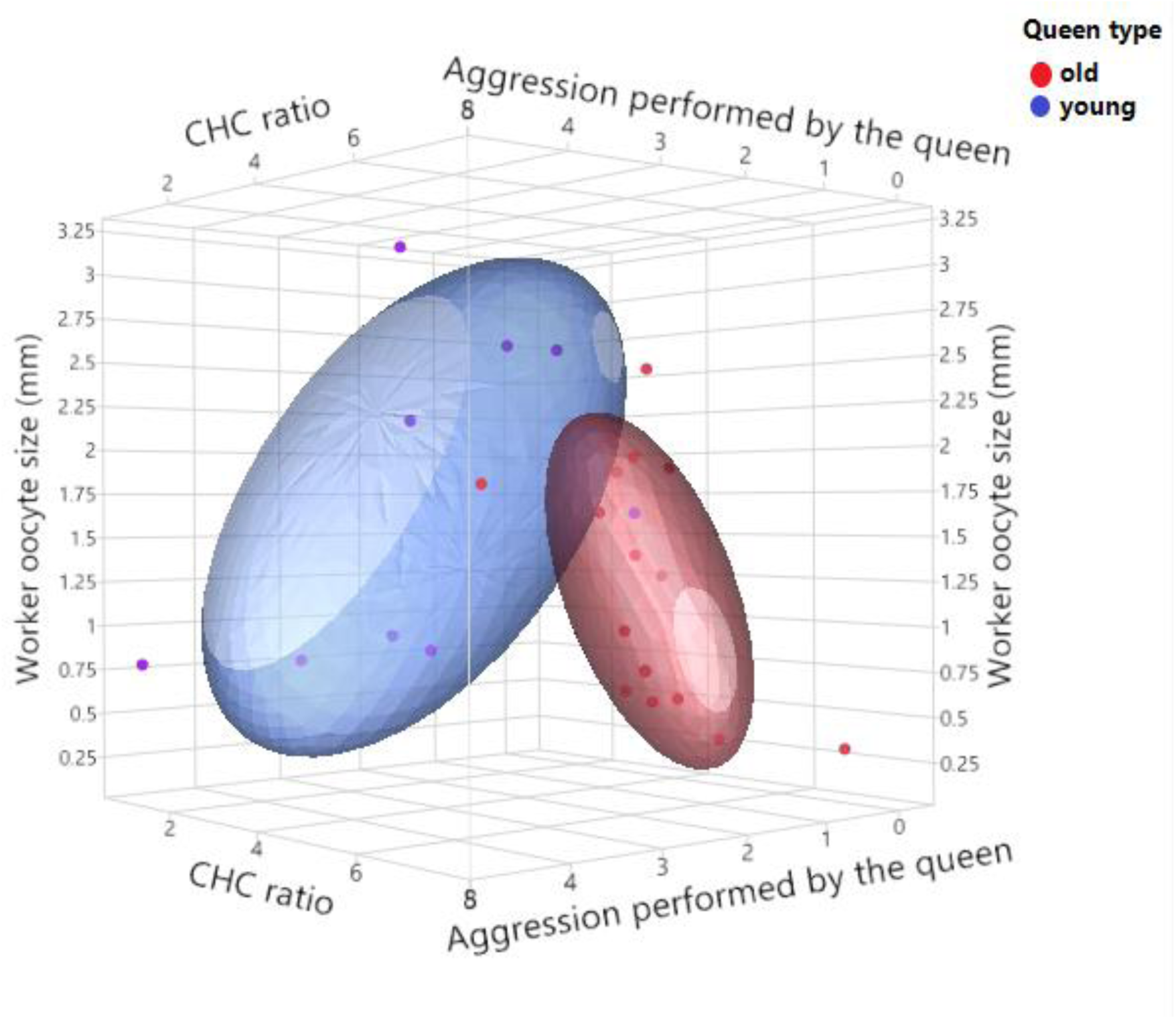
Aggression and short to long CHC ratio in queens as predictors for worker oocyte size in groups headed by young and old laying queens. Different queen types are indicated by different colours. Confidence ellipsoids indicate the distribution of data in each group.

## Discussion

Our results indicate significant changes occurring in the behaviour and chemical profile of *Bombus impatiens* queens, and in their capacity to inhibit worker reproduction, as they progress through their life cycle. Our findings suggest that the mechanism – and the efficacy – of reproductive regulation in *Bombus impatiens* changes with the queen’s age and her life stage. Newly mated and young egg-laying queens rely on aggression to inhibit worker reproduction. This strategy is likely effective in young egg-laying queens since the queen’s aggressive behaviour was the only significant negative predictor of ovarian activation in workers. However, in old egg-laying queens that were most effective at inhibiting worker reproduction while performing the least aggressive behaviour, this is clearly not the case. Old egg-laying queens’ cuticular chemistry, but not aggressive behaviour, was a significant negative predictor of worker ovary size. These findings indicate that as the queens age, they change their dominance strategy, shifting from aggression towards chemical signalling as would be adaptive under the changing social conditions: a young egg-laying queen would be confronted by a small number of workers but an old egg-laying queens would face hundreds of potential adversaries.

Both types of egg-laying queens, and only them, caused meaningful reduction in worker oocyte size. These queens supposedly provide indirect fitness benefits for workers, suggesting that workers were able to perceive the quality of the queen, and that her dominance signal is apparently honest. This honest signal is probably the egg-laying behaviour combined with either aggressiveness in young egg-laying queens or with S:L CHC ratio in old egg-laying queens. S:L CHC ratio increased steadily as function of age, paralleling the increase in colony productivity and fitness value to workers, peaking in production of sexuals. It should be noted however that although queen and workers in our study were unrelated, to our knowledge, there are no evidence in bumble bees that workers are capable to recognize kin from non kin (Amsalem and Hefetz, 2010) and thus we assume that workers perceive the queen as their mother.

Queen types differed not only in their CHC profile, but also in the proportions of long chain acetates (higher in young egg-laying queens) and aldehydes (higher in unmated queens). Acetates were found in other bee species, but their biological significance is still poorly understood (Cane, 1983; Hefetz et al., 1996; Hefetz et al., 1993). Aldehydes in unmated queens might serve for sexual communication, but also, being an intermediate product in hydrocarbon biosynthesis (Blomquist et al., 2010), might simply indicate high metabolic activity in newly emerged queens. How different components of the cuticular profile might be linked to a queen’s productivity and whether CHCs in lab-reared colonies differ from wild colonies is yet unknown and future research on the subject is needed.

The change in CHC ratio between young and old egg-laying queens is likely driven by intrinsic factors. In our study, young egg-laying queens, although maintained in the same conditions throughout the experiment, changed their cuticular chemistry gradually as they aged. This finding alludes that the queen’s plasticity in behaviour and signalling and the ability to change the dominance strategy evolved as an adaptation to changing social environment during their lifetimes, i.e., that queens evolved to be flexible. A somewhat similar phenomenon was documented in ponerine ants where the alpha female aggressiveness and CHC profile gradually change following mating (Monnin et al., 2002) and was proposed for domestic hens where focal individuals displayed less aggressive behaviour as a function of increasing group size (Estevez et al., 2003; Pagel and Dawkins, 1997). More research, however, is required to elucidate the degree of plasticity in behaviour and cuticular chemistry that a queen can exhibit in different social situations.

The most prominent difference found between the cuticular profile of old egg-laying queens and other queens was in the relative amounts of short (≤24 carbons) and long (≥26 carbons) alkanes and alkenes that changes with age in different queen types. Cuticular hydrocarbons are ancient and highly conserved semiochemicals serving for a variety of communicative purposes, such as sexual signalling, nestmate recognition, and royal status signalling (Blomquist and Bagnères, 2010; Smith and Liebig, 2017). These findings harken back to studies that found a similar age dependent (and fertility-dependent) change in the cuticular composition of *Musca domestica* (Mpuru et al., 2001). Thers, a similar shift in abundance of chain lengths longer and shorter than 25 carbons is driven by differential expression of chain-length-specific biosynthetic genes, which, in turn are regulated by ovarian maturation (Vaz et al., 1988). The change in the hydrocarbon profile in our study is probably not driven by ovarian activation as both young and old egg-laying queens possessed fully developed ovaries and were actively laying eggs. However, it is possible that conserved biosynthetic mechanisms, such as chain-length specific elongase enzymes, were co-opted for the new, social environment and controlled by novel regulatory means (Robinson and Ben-Shahar, 2002).

Overall, it is the change in the overall chemical profile rather than the quantitative difference in any single compound that accounts for differences between the queen types and is important for maintaining the queen’s reproductive monopoly. Previous studies suggested that one or another single hydrocarbon played the role of a queen pheromone in social insects (Oliveira et al., 2016; Van Oystaeyen et al., 2014), but based on our findings this doesn’t seem to be the case. Amounts of pentacosane, previously suggested to regulate reproduction in *B. terrestris* (Van Oystaeyen et al., 2014), did not show any consistent differences between queens in our study and were not a significant predictor of worker oocyte size. Actually, it was as high in unmated queens that were incapable of reducing worker reproduction as it was in old egg-laying queens that were most effective in doing so. These results corroborate previous findings that showed the importance of the complete chemical profile for the perception of a fertility signal (Smith et al., 2015)

It is important to note that aggression among workers was also a significant predictor of oocyte size in workers: worker ovary size increased with increasing aggression among workers, but only in groups headed by unmated and old egg-laying queens (with much higher aggression in workers paired with unmated queens), where the queen herself performed very little aggression. This finding can be interpreted in two ways: either that workers with higher reproductive potential are more actively vying for dominance, or that aggressive competition and assertion of the dominant status are necessary for workers to develop their ovaries, as was shown in wasps (Lamba et al., 2007; Toth et al., 2014). The latter is better supported by our data, since aggression was a significant predictor of worker ovaries only in groups where the queen performed little dominance behaviour. The contribution of workers dominance behaviour to reproductive partitioning and its interplay with the queen’s efforts to monopolize reproduction is a subject that merits further study.

Overall, our results present an interesting insight into the mechanisms underlying reproductive dominance in social animals and suggest that these mechanisms are more complex and flexible than previously thought with substantial differences across different life stages, characterized by a shift between aggressive behaviour to chemical signalling and age-driven increase in S:L CHC ratio within the same species.

